# Automated wildlife re-identification by merging information from multiple body parts: A case study in sea turtles

**DOI:** 10.64898/2026.08.28.747856

**Authors:** Lukáš Adam, Micol Montagna, Valeria Roma, Agnese Mancini, Kostas Papafitsoros

## Abstract

Wildlife re-identification (re-ID) is a widely used and powerful tool with diverse applications in animal ecology and conservation. Current automated methods typically operate on single images of a single body part of the animal. However, a single encounter may contain multiple images capturing different body regions, each providing complementary individual-specific information. In contrast to automated approaches, researchers often manually select the most suitable images and regions for identification based on factors like visibility, occlusion and image quality. This creates a mismatch between automated methods and field practice, limiting the practical adoption of current automated re-ID pipelines. Here, we address this by introducing an encounter-based, multi-body-part re-ID framework, using sea turtles as a model taxon. Our framework combines three elements: (1) An orientation-aware deep learning model, TurtleDetector, that in addition to the full bodies, it also automatically segments key body regions, i.e. heads, front and hind flippers, from images within an encounter; (2) a hybrid body-part-specific retrieval method, that sequentially combines a fast global-feature model (MiewID or DINOv3) with a more accurate but costlier local-feature model (ALIKED with LightGlue); and (3) a merged identity-prediction strategy that selects the highest calibrated similarity score across all available body parts and images of an encounter. We evaluate the framework on three long-term re-ID datasets spanning three species, loggerheads, greens, and hawksbill turtles, under an evaluation protocol that mirrors real-world, time-aware re-ID workflows. Across datasets, combining multiple body regions consistently improved identification performance over the best-performing single body region, resulting to an increase of 4–6% in top-1 accuracy. Interestingly, body regions traditionally underused in sea turtle re-ID, such as the hind flippers and carapaces, provided complementary identifying information that improved encounter-level re-ID when integrated through the hybrid retrieval method. Our findings demonstrate that automated wildlife re-ID can benefit from moving beyond single-image, single-body-part identification towards encounter-level integration of all available visual evidence. Our work further suggests that, where feasible, field photo-acquisition protocols should aim to capture multiple informative views of an individual during each encounter. Importantly, many species and taxa, including elephants, primates, cetaceans, and other large vertebrates, possess such individual-specific features across multiple body regions, highlighting the broad potential applicability of our framework.

## 1. Introduction

Wildlife re-identification (re-ID) denotes the task of identifying individual animals from images acquired across space and time (Vidal *et al*., 2021). It provides a non-invasive analogue of traditional capture-mark-recapture methods, enabling large-scale individual monitoring without physical tagging and thereby avoiding animal stress, potential injuries, and associated logistical challenges. The identification is performed by exploiting any external morphological characteristics, features, patterns that are both stable over the timescale of the study (and often permanent) and unique to each individual. Examples of such patterns are stripes, spots, scales, skin/fur patterns, body shapes and contours, just to name a few. Wildlife re-ID is a well-established and indispensable tool in ecological research, enabling studies that would otherwise be impossible to conduct, with many relevant projects operating for decades. Such studies include inferring population abundance and density (Parham *et al*., 2017), estimating individual survival rates (Schofield *et al*., 2020), studying social behaviour (Bond *et al*., 2021), quantifying disease and anthropogenic pressures (Papafitsoros *et al*., 2021), and several others.

Inevitably, the ever-increasing volume of imaging data generated by such studies, combined with limited human resources, creates a significant bottleneck, as manually inspecting often tens of thousands of images becomes infeasible. To address this challenge, numerous automated re-ID methods and pipelines have been developed during the last 15 years, greatly accelerating the comparison of large numbers of images. We refer the reader to the comprehensive reviews by Kühl and Burghardt, 2013; Vidal *et al*., 2021; Stewart *et al*., 2021; Čermák *et al*., 2024 for an overview of the field, its historical development, and the categorisation of existing methods. Such methods operate under the framework of image retrieval, where new *query images* (unidentified individuals) are compared with images of a *database* (previously identified individuals) and the most similar images are retrieved for a final inspection and identification by a human, thus sparing the expert from searching the whole database. Image similarity is either inferred by comparing deep feature vectors (*embeddings*), extracted by deep neural networks (deep metric learning), e.g. MegaDescriptor and MiewID (Čermák *et al*., 2024; Otarashvili *et al*., 2024) or by locating and comparing common local features (*keypoints*) that appear in image pairs (Lowe, 2004; DeTone *et al*., 2018; Zhao *et al*., 2023), or by combining these two techniques (Cermak *et al*., 2024); we refer to Section 2 for more details.

Despite the tremendous progress of the field, which is largely driven by advances in machine learning and computer vision and the ever-increasing availability of training data, the practical adoption of automated re-ID methods by ecologists and biologists has lagged behind, with only a small fraction of projects benefiting from these developments (Picek *et al*., 2026a). The practical deployment of these methods is hindered, among other factors, by discrepancies between how the animal re-ID task is formulated and evaluated in computer science and how it is applied in real-world ecological settings. One particular instance of this discrepancy is the use of multiple body parts during the identification process. In many animal species, unique identifying features can be found in several parts of their bodies, which are not always all visible in a single image. Examples include features located at the left and right sides of the body (Parham *et al*., 2017; Adam *et al*., 2025), heads and flippers in sea turtles (Papafitsoros *et al*., 2025), tail flukes and fins in cetaceans (Patton *et al*., 2023), tusks and ears in elephants (Kulits *et al*., 2021) etc. Ecologists typically examine *sets of images of the same individual* collected during a *single encounter*, selecting the most informative body parts for identification based on factors such as image quality and the absence of occlusions, and then attempt to match accordingly. If one body part does not yield a match, alternative body parts may still provide sufficient information; thus, the search procedure is often repeated using them. In practice, therefore, animal re-ID is inherently *multi-body part* and *encounter-based*, not single image-based. In contrast, automated pipelines usually compare single images, focusing on a single body part against a database. This study addresses this discrepancy by introducing an automated re-ID framework that leverages information from multiple body parts, using sea turtles as a model taxon.

### Our contribution

We propose a general re-ID framework for ecological monitoring that operates at the level of encounters and across multiple body-part information. The framework leverages multiple body regions for identification and performs identity inference by using complementary information across regions extracted from multiple images of single encounters; see Figure 1 for a visualisation. While we use sea turtles as model species, the framework is applicable to other taxa exhibiting identifying patterns in different body regions. In particular, we make the following contributions:

i. **Orientation-aware multi-body part detection enabling encounter-level inference:** We introduce *TurtleDetector* ^1^, a deep learning-based model for automatic segmentation of multiple identifying body regions (full body including carapace, head, front and hind flippers, and left/right sides) across images within an encounter. The model successfully segments multiple body parts in an orientation-aware manner, across all three sea turtle species, with about 90% of encounters yielding at least five extracted body parts.
ii. **Fast and accurate hybrid body part-specific matching:** For each body part, we propose a hybrid matching model via the sequential combination of two state-of-the-art re-ID models: A deep metric learning-based feature extractor (we use MiewID or DINOv3), which is computationally efficient but less accurate and a local feature extractor (ALIKED with LightGlue), which is more accurate but computationally expensive. We show that their combination yields an overall improvement in the accuracy-efficiency trade-off compared to each model alone, leading to scalable and accurate body-part specific matching.
iii. **Identity prediction based on merged calibrated multi-body part similarity score strategy:** In order to perform matching (inference) exploiting information from multiple body parts, we propose a novel merged similarity score strategy which uses the largest similarity scores from each body part after these have been calibrated. Under this strategy, the identity prediction for an encounter is determined by the most informative body region across all encounter images.
iv. **Evaluation under realistic re-ID workflows and multiple datasets:** We evaluate the proposed framework by performing experiments that are aligned with the real-world re-ID workflow: Multi-body part information from a given encounter is compared to multi-body part information from all the previous encounters. We do so using two curated datasets of three sea turtle species (one from Egypt, and one from Greece).
v. **Multi-body-part re-ID outperforms single-body-part approaches:** Our results imply that using the hybrid matching model and multiple body part information leads to a gain of up to 6% Rank-1 accuracy over the standard single body part approaches. As additional species-specific contributions, we demonstrate the feasibility of hind-flipper-based re-ID across all three sea turtle species and of carapace-based re-ID in green turtles.
vi. **Open-source code and models:** We show that it is possible to analyse 9 years of data within hours on a CPU with over 90% top-1 accuracy. We hope that, together with making the code and the TurtleDetector model open-source, this will motivate other researchers to use our tools.

**Figure 1.**
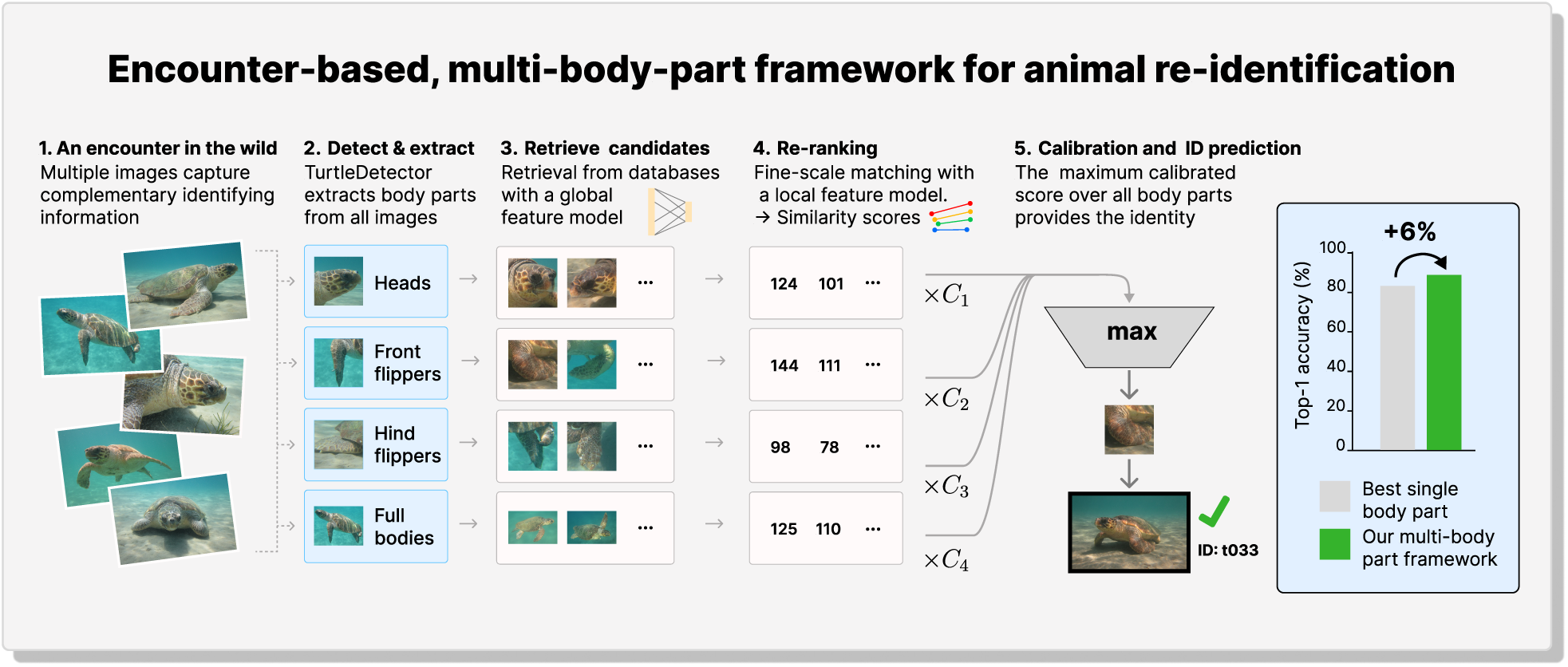
Visualisation of the proposed encounter-based, multi-body-part framework for animal re-identification.

## 2. Overview of wildlife re-ID methods

### 2.1. State-of-the-art approaches

State-of-the-art re-ID methods can be broadly divided into two categories: Global feature and local feature approaches, see Figure 2. Global feature re-ID methods are based on deep metric learning and leverage deep neural networks to represent each image with a deep feature vector (embedding), e.g., MegaDescriptor (Čermák *et al*., 2024) and MiewID (Otarashvili *et al*., 2024). During training, the networks are optimised such that images of the same individual are mapped to nearby points in the embedding space, whereas images of different individuals are mapped further apart, typically using cosine similarity as the distance metric. These methods are trained on large curated datasets, and their inference is based not only on identifying characteristics from geometric patterns but also on richer ones such as colouration and texture. Once trained, they are highly computationally efficient at inference, as identification reduces to computing pairwise similarities between feature vectors, a simple and scalable operation.

**Figure 2.**
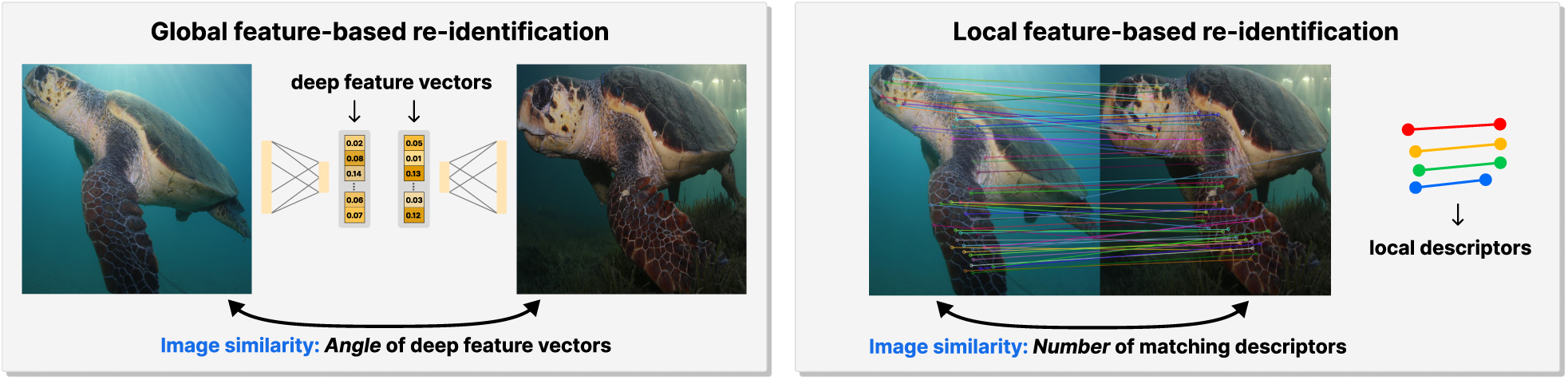
Global versus local feature methods for animal re-ID. In global feature-based re-ID, image similarity is measured by comparing the alignment (angle) of feature vectors extracted by a deep neural network. The latter has been trained so that images of the same individual animal produce more closely aligned feature vectors than images of different individuals. In contrast, local feature-based methods identify salient local features in each image and assess similarity by matching corresponding local descriptors across an image pair, with the number of successful matches serving as a measure of similarity.

On the other hand, modern local feature methods, such as Superpoint (DeTone *et al*., 2018) and ALIKED (Zhao *et al*., 2023), detect salient local image features and represent them using local descriptors. Identification is then performed by matching these descriptors between a query image (unknown identity) and database images (known identities), with the number of matching descriptors serving as a similarity score. These methods are very effective in detecting geometric similarities and therefore well-suited for patterned species, including sea turtles (Adam *et al*., 2025). However, unlike global feature approaches, their computational cost scales poorly with database size, as descriptor matching must be performed for every query-database image pair and is substantially more expensive than comparing feature vectors.

All re-ID methods essentially operate under the framework of image retrieval. Similarity scores are computed between the query image and each image in the database. Then, the *k* most similar database images to the query are retrieved for final inspection by an expert, with *k* usually small. The top-*k* accuracy is naturally chosen to evaluate the method, measuring the proportion of query images with at least one correct identity prediction among the *k* retrieved images. This reflects the common ecological workflow in which researchers inspect a small number of the highest-ranked matches rather than relying solely on the top prediction.

### 2.2. Sea turtle re-ID

Sea turtle re-ID is predominantly based on the polygonal scales of the head, primarily using the left and right facial profiles and, less frequently, the dorsal (top) view of the head (Schofield *et al*., 2008; Carpentier *et al*., 2016; Papafitsoros *et al*., 2025). Consequently, most automated workflows have focused on head images (Dunbar *et al*., 2014; Dunbar *et al*., 2021; Adam *et al*., 2025; Polychronou *et al*., 2026), and state-of-the-art deep feature extractors are trained predominantly with images of this body region (Otarashvili *et al*., 2024; Čermák *et al*., 2024).

Nevertheless, the scale patterns of the front flippers have also been proposed as a valuable source of identifying information (Caillouet Jr *et al*., 1989; Pursley, 2020; Papafitsoros *et al*., 2025). Indeed, an upcoming chapter of the Research and Management Techniques for the Conservation of Sea Turtles, by IUCN/SSC Marine Turtle Specialist Group (Papafitsoros *et al*., 2025) states that “*Thus, while we still recommend that sea turtle photo-ID be based on facial scales, flipper scales may be a valuable secondary resource, especially when combined with head scales, and it has been shown to enhance* (manual) *matching accuracy in practice.*”. In practice, however, manual inspection of the flippers is considerably more laborious than inspection of facial scales because the scales are smaller and often more difficult to discern. As a result, only a handful of automated approaches have explored flipper-based identification, typically under controlled conditions (Pursley, 2020; Mills *et al*., 2023). Furthermore, the potential of hind flipper scales for sea turtle re-ID has never been investigated, despite this often being the only visible identifying region in skittish individuals that rapidly swim away from the photographer (Papafitsoros *et al*., 2025).

Finally, carapace-based sea turtle re-ID has generally been regarded as the least promising approach for sea turtles and has therefore received comparatively little attention (Tabuki *et al*., 2021). One reason is that the carapace is often covered by algae and other epibionts, obscuring its underlying pigmentation pattern. In addition, manual comparison of carapaces is considerably more tedious and time-consuming than comparison of facial scale patterns. Perhaps most importantly, the pigmentation pattern of the carapace is not always temporally stable (Tabuki *et al*., 2021; Papafitsoros *et al*., 2025), limiting its suitability as a standalone identifying feature.

These characteristics make sea turtles an ideal model species for the development and testing of automated re-ID frameworks that integrate information from multiple body parts and operate at the encounter level rather than on individual images alone.

## 3. Methodology

### 3.1. Testing datasets

We use two datasets, SeaTurtleID2022 and TurtlewatchEgypt, corresponding to three different sea turtle species: loggerhead sea turtles (*Caretta caretta*) for the former, and greens (*Chelonia mydas*) and hawksbills (*Eretmochelys imbricata*) for the latter.

The SeaTurtleID2022 dataset (Adam *et al*., 2024) includes underwater images of loggerhead sea turtles taken on Zakynthos island, Greece. The dataset comprises 8,729 images of 438 unique individuals recorded across 1,221 encounters between 2010 and 2022. We refer the reader to Adam *et al*., 2024 and Schofield *et al*., 2020 for details on data collection. The TurtlewatchEgypt dataset contains underwater images of green and hawksbill sea turtles. It originates from the Southern Egyptian Red Sea and was compiled by the TurtleWatch Egypt 2.0 project, managed by the NGOs Marine Life Watch and Marine Life Conservation and Preservation Foundation (Montagna *et al*., 2017; Montagna *et al*., 2023; Mancini *et al*., 2025). It is part of a larger collection project that includes citizen science photos. For the numerical experiments in this paper, we selected a subset collected by the core members of TurtleWatch Egypt 2.0, comprising 2,058 encounters of green and 126 of hawksbill sea turtles, collected between 2017 and 2025, corresponding to 321 uniquely identified individuals. All images were acquired during underwater surveys conducted by trained researchers. Individual identities were established through manual comparison of photographs with the catalogue of previously identified turtles, resulting in a fully identity-annotated dataset.

For both datasets, whenever possible, multiple photographs of the same individual were taken during a single encounter to maximise subsequent identification success. Priority was given to capturing the left and right facial profiles, complemented by images of the whole body and other distinctive natural markings, including injuries, scars, epibionts, and other individual-specific characteristics. Although flippers were not originally considered a primary identification feature and were therefore not systematically targeted, they were frequently captured during head-profile photography.

Figure 3 shows sample images of the two datasets, and Figure 4 shows the number of encounters per year for SeaTurtleID2022 (left) and TurtlewatchEgypt (right). Relevant to our later experiments, we also show the number of “known” (orange) and “new” (green) encounters for each dataset. The former contains individuals that had already been observed in a previous year, whereas the latter contains new individuals.

**Figure 3.**
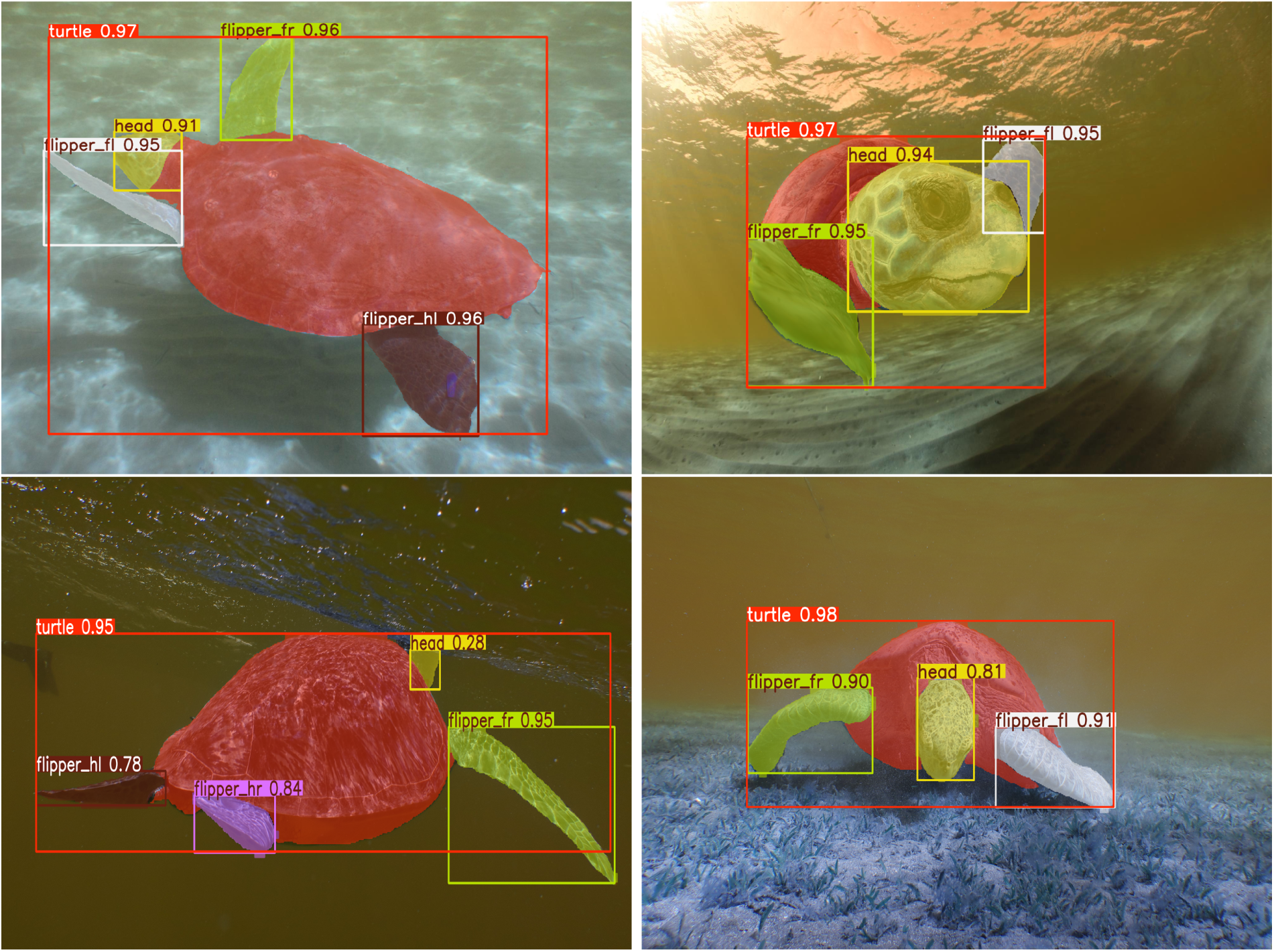
Sample images of SeaTurtleID2022 (top) and TurtlewatchEgypt (bottom) datasets. Here, we also include the segmentation results of TurtleDetector showing the categories “turtle” (full body), “head”, “flipper fl” (front left), “flipper fr” (front right), “flipper hl” (hind left) and “flipper hr” (hind right). The numbers show the confidence of the segmentation masks.

**Figure 4.**
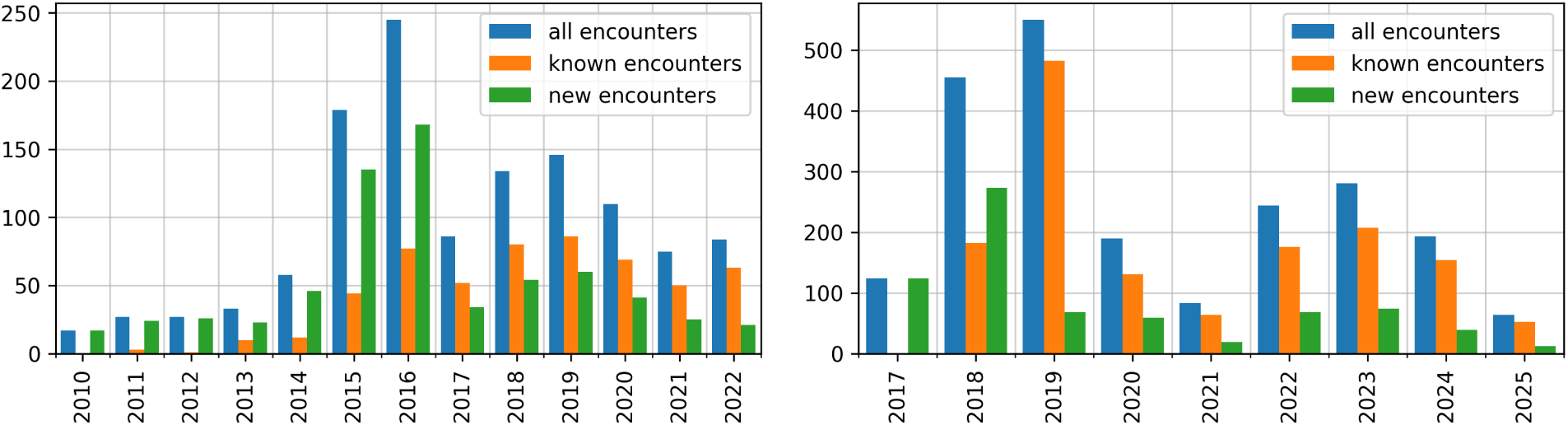
The total number of encounters per year (blue) for SeaTurtleID2022 (left) and TurtlewatchEgypt (right). The encounters are divided, depending on whether the individual was known (orange) or new (green), i.e. whether it had been observed or not in one of the previous years, respectively.

### 3.2. Image retrieval experiments towards a realistic time-aware re-ID workflow

To emulate a realistic re-ID workflow, we adopt a time-aware, encounter-based evaluation protocol based on consecutive years of data collection. Rather than performing a single database-query split, we conduct a series of “annual” retrieval experiments. For each study year, all photographs from previous years constituted the reference database, while photographs collected during the current year formed the query set. Consequently, the reference database progressively expanded over time, reflecting the accumulation of identified individuals in a real monitoring program. We remark that this evaluation protocol differs from standard practices in the computer vision literature, where image retrieval methods are commonly evaluated using an image-based random split protocol (Adam *et al*., 2024; Picek *et al*., 2026a). This would be here unsuitable since, under a random split, images from the same encounter may appear in both sets, making retrieval substantially easier and inflating performance.

We also adopt an encounter-aware evaluation, again reflecting the real-world practice. Rather than assigning an identity to individual images, the objective here is to assign a single identity to an entire encounter. This requires localising the body regions across all images from an encounter that provide the most distinctive visual information for identification. For example, although sea turtles are most commonly identified by their facial scale patterns, an encounter may contain only images in which the head is poorly visible (see the bottom-left photo in Figure 3). In such cases, identification may be more reliably achieved using, for instance, a photograph with a clearly visible front or hind flipper.

In this work, we are mainly interested in comparing the identification performance (discriminatory capabilities) of the proposed multi-body-part, encounter-based framework with that of conventional single-body-part approaches. It is thus sufficient to perform this comparison in a closed-set image retrieval setting, where each query image corresponds to an individual already present in the reference database. Thus, new encounters, i.e. green columns in Figure 4, are not included in the query images but retained in the database. For example, if an individual is first encountered in 2019, that encounter is not used as a query in the 2019 evaluation but is included in the reference databases for 2017-2019, 2017-2020, and all subsequent evaluation periods. This brought the total number of query encounters to 547 and 1,448 for SeaTurtleID2022 and TurtlewatchEgypt, respectively.

The retrieval performance was evaluated using top-1 and top-*k* accuracy adapted for encounter-based identification. Thus, top-1 accuracy measures the proportion of query encounters for which the correct individual is ranked first. Top-*k* accuracy measures the proportion of query encounters for which the correct individual appears among the first *k* retrieved candidates. Performance was computed independently for each yearly retrieval experiment and then averaged across all evaluation years to obtain the final reported results.

### 3.3. Segmenting body parts – TurtleDetector

To exploit the complementary information present in sea turtle heads, flippers and carapace, we developed *TurtleDetector*, a novel deep learning model for automatic body-part detection and segmentation. Given an input image, TurtleDetector localises the full body and all visible identifying body parts, and for each, predicts a bounding box, a segmentation mask, and an orientation-aware body-part label. The predicted labels distinguish the head and the four flippers: front left (FL), front right (FR), hind left (HL), and hind right (HR). Each prediction is further associated with a confidence score in the range [0, 1], indicating the model’s confidence in the localisation and classification of the detected body part, see Figure 3.

TurtleDetector was trained in two stages. In the first stage, the model was trained on the existing orientation-aware body-part segmentation annotations from the SeaTurtleID2022 dataset. The second stage fine-tuned it on terrestrial sea turtle images from the TurtlesOfSMSRC project to improve generalisation across acquisition environments. Since body-part annotations were not available for the latter images, we created them as part of this work. Segmentation masks for heads and flippers were generated using SAM3 (Carion *et al*., 2026) while orientation-aware body-part labels were assigned using a semi-automatic heuristic procedure. The complete annotation pipeline, together with the training scripts for TurtleDetector, is publicly available on the project’s website. The resulting detector accurately localises and segments the full body and individual turtle body parts in both underwater and terrestrial images across multiple sea turtle species.

### 3.4. Identity prediction for encounters

We now describe our complete pipeline for identity prediction. The pipeline operates on encounters and consists of four stages: (i) body-part extraction, (ii) computation of similarity scores between the query encounter and the database, (iii) scaling the similarity scores, and (iv) identity assignment based on these scores.

The first step is to extract the identifying body parts from every image in an encounter and assign them orientation-aware labels. The SeaTurtleID2022 dataset already provides body-part annotations, whereas for TurtlewatchEgypt, they are obtained automatically using TurtleDetector. To reduce the computational intensity, for each TurtlewatchEgypt encounter, we retain at most 10 high-quality detections, ranked by their confidence scores. By default, only detections with a confidence score greater than 0.8 are considered. If fewer than two detections are available for a particular body part, the confidence threshold is decreased. For SeaTurtleID2022, all body-part annotations are used since they represent the ground truth. In addition to the individual body parts (head and flippers), we also include full-body images in our experiments. This serves two purposes. First, full-body images contain the carapace, allowing us to investigate whether incorporating identifying information from this additional body region further improves the proposed multi-body-part framework. Second, because full-body images encompass all visible body parts, they provide a natural baseline for assessing the value of explicit body-part localisation. In particular, they enable us to evaluate whether detecting individual body parts and matching them separately yields more accurate identification than applying re-ID directly to the full-body image. In what follows, we consider that the encounter is represented by 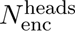 head images, 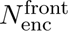 front-flipper images, 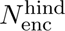 hind-flipper images and 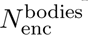 full body images and that the database consists of 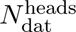 head images, 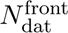 front-flipper images, and 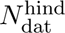 hind-flipper images and 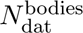 full body images.

The second step computes the similarities between the query encounter and the reference database using one of two alternative retrieval strategies, which we describe next. The first strategy uses only a global feature method. A global embedding is extracted for each body-part image and compared with those of all database images of the same body part. Since only the same body parts are compared, this amounts to the computation of

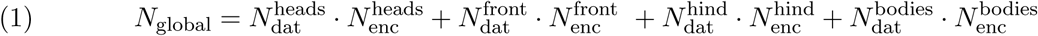

*global similarity* scores. The second strategy combines a global with a local feature model, see also Cermak *et al*. (2024), and Yesharim *et al*. (2026); Algasov *et al*. (2026) for a similar approach. It first computes the same global similarity scores as in the first strategy. These similarities are then used to retrieve a *Budget* of *B* most similar database images for each query body part, where *B* is substantially smaller than the total number of database images. Local-feature matching is then performed only for these candidate pairs, producing refined similarity scores that re-rank the initial retrieval results. This procedure, in addition to (1), requires the computation of

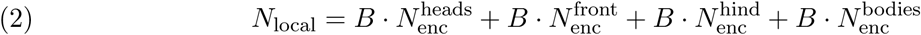

*local similarity* scores.

The third step takes the *N*_global_ or *N*_local_ scores computed in the previous step and scales each body part individually. We use the simplest scaling possible. For the global features, we do not use any scaling because the similarity scores are in the same range [−1, 1]. For the local features, the similarity score is the number of matching keypoints. Since heads typically contain fewer distinctive visual patterns than flippers (fewer scales), their similarity scores are typically smaller, and we multiply them by a calibration constant *C*_calibration_ ≥ 1.

The fourth step aggregates the, either *N*_global_ or *N*_local_, similarity scores across all body parts and orders the corresponding database images from the highest similarity to the lowest. The ranked list is then converted into an ordered list of candidate identities by retaining only the highest-ranked occurrence of each individual, ensuring that every identity appears at most once. The first identity is used for top-1 evaluation, while the first *k* identities are used for top-*k* evaluation.

### 3.5. Orientation-aware vs orientation-unaware matching for flippers

To fully leverage the orientation-aware body-part labels provided by the TurtleDetector, we incorporate orientation awareness for flippers and compare only flippers with the same orientation, that is, front left with front left flippers and similarly for other orientations. The improvement achieved by this strategy depends on the quality of the orientation labels. If the orientation labels are correct, the strategy restricts the comparisons to anatomically corresponding body parts, which should improve the performance. On the other hand, incorrect orientation labels may prevent the correct match from being considered, potentially reducing the retrieval performance.

### 3.6. Implementation details

We use ALIKED as the local feature model. For TurtlewatchEgypt, we use MiewID (Otarashvili *et al*., 2024) as the global feature model, as it has been shown to perform well across these species. For SeaTurtleID2022, however, we could not use either MiewID or MegaDescriptor because this dataset was part of the training data for both models, which would lead to inflated performance. Instead, we use DINOv3 (Siméoni *et al*., 2025) as the global-feature model for candidate retrieval. For the calibration constant for the ALIKED similarity scores, after some experimentation, we selected *C*_calibration_ = 1.25 as the similarity score multiplier for the heads for TurtlewatchEgypt and *C*_calibration_ = 1 for SeaTurtleID2022. In Table 1, we summarise the similarities and differences between the retrieval pipeline components used for the two datasets.

**Table 1.** Similarities and differences between the retrieval pipeline components used for the two datasets.

|  | SeaTurtleID2022 | TurtlewatchEgypt |
| --- | --- | --- |
| Species | Loggerheads | Greens & Hawksbills |
| Origin | Zakynthos, Greece | Red Sea, Egypt |
| Number of images | 8,729 | 32,512 |
| Number of encounters | 1,221 | 2,184 |
| Number of individuals | 438 | 321 |
| Year span | 2010-2022 | 2017-2025 |
| Segmentation masks | Present in the dataset | Generated by TurtleDetector |
| Number of masks per body part | All | 0-10 |
| Global extractor | DINOv3 | MiewID |
| Local extractor | ALIKED | ALIKED |
| $C_{\text{calibration}}$ | 1 | 1.25 |

### 3.7. Running times

This section presents the running times for the individual components of the pipeline. We show the times on both a GPU (NVIDIA GeForce RTX 3070) and a CPU employed in facilities without specialised computing infrastructure. The presented times should be considered approximate due to issues such as parallelisation and the ambiguity on the definition of tasks (e.g. whether one considers image transform being part of image loading or of running TurtleDetector).

Table 2 shows the running times for TurtleDetector in milliseconds when processing a single image. Image loading is significantly slower for TurtlewatchEgypt because the dataset contains original images of high resolution, unlike SeaTurtleID2022, where most images were resized to 2000 × 1333 pixels. The difference in the running times for TurtleDetector is caused by the need to transform the segmentation masks back to the larger photos in TurtlewatchEgypt. Even in the worst case (TurtleWatch + CPU), TurtleDetector can process over 10,000 images per hour, while this number can be ten times as high for the best case (SeaTurtleID2022 + GPU).

**Table 2.**
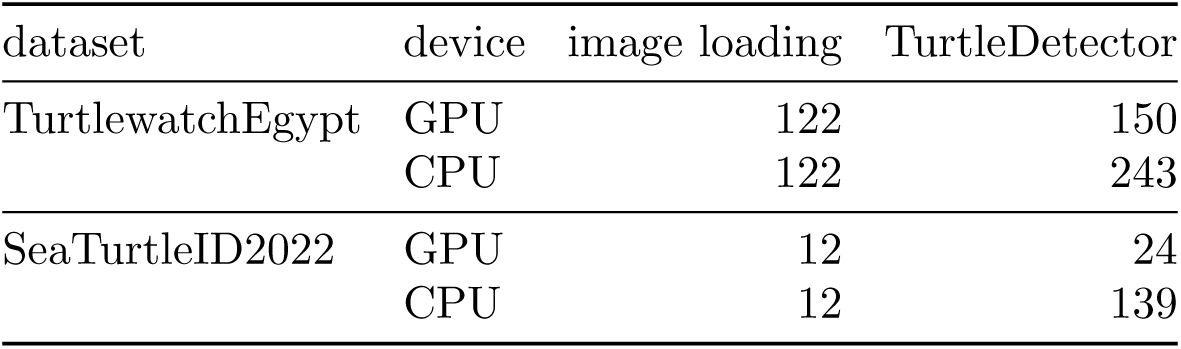
Running times for TurtleDetector (rounded in milliseconds).

| dataset | device | image loading | TurtleDetector |
| --- | --- | --- | --- |
| TurtlewatchEgypt | GPU | 122 | 150 |
|  | CPU | 122 | 243 |
| SeaTurtleID2022 | GPU | 12 | 24 |
|  | CPU | 12 | 139 |

Table 3 shows the running times for individual components of the pipeline in milliseconds when processing a single extracted body part. Even though, as expected, the CPU is significantly slower than the GPU, this difference becomes less relevant for TurtlewatchEgypt due to the large image loading times. The important part is the feature matching (similarity computation) for MiewID and DINO, which is essentially zero when rounded to the nearest millisecond. This explains why it is computationally feasible to consider the entire database when computing the *N*_global_ global similarity scores (1). Since the feature matching is more expensive for the local-feature method ALIKED, it is preferable not to be performed across the whole database, but only for the budget *B* as specified in (2).

**Table 3.**
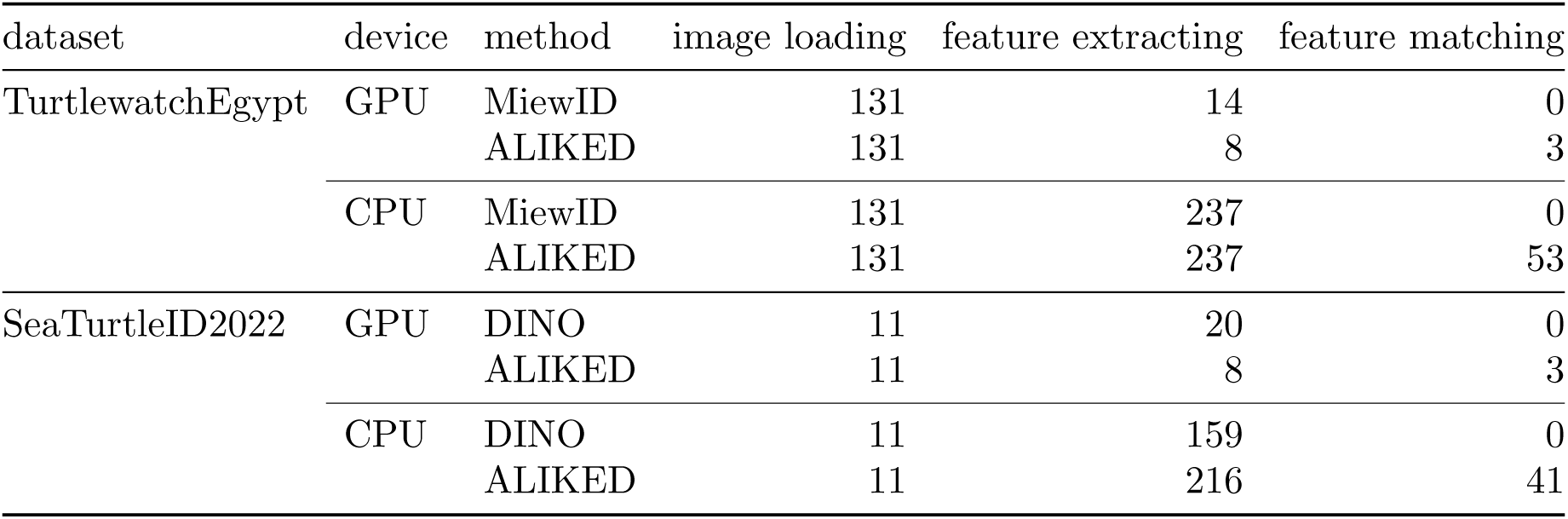
Running times for individual components of the pipeline (rounded in milliseconds).

| dataset | device | method | image loading | feature extracting | feature matching |
| --- | --- | --- | --- | --- | --- |
| TurtlewatchEgypt | GPU | MiewID | 131 | 14 | 0 |
|  |  | ALIKED | 131 | 8 | 3 |
|  | CPU | MiewID | 131 | 237 | 0 |
|  |  | ALIKED | 131 | 237 | 53 |
| SeaTurtleID2022 | GPU | DINO | 11 | 20 | 0 |
|  |  | ALIKED | 11 | 8 | 3 |
|  | CPU | DINO | 11 | 159 | 0 |
|  |  | ALIKED | 11 | 216 | 41 |

Based on the information of Table 3, we can also compute the number of images that can be processed in one hour, see Figure 5. Even with the slowest CPU computation for TurtlewatchEgypt, we can process 3,200 images per hour for *B* = 10 under our experimental design, and this number drops to 600 for *B* = 100. This showcases the high efficiency of our framework. For instance, the full database of the TurtleWatch Egypt 2.0 project (not used is this work in its entirety) contains approximately 32,000 images. This means that the proposed pipeline with *B* = 10 enables predicting identities for 9 years of data collection in just a few hours without a GPU.

**Figure 5.**
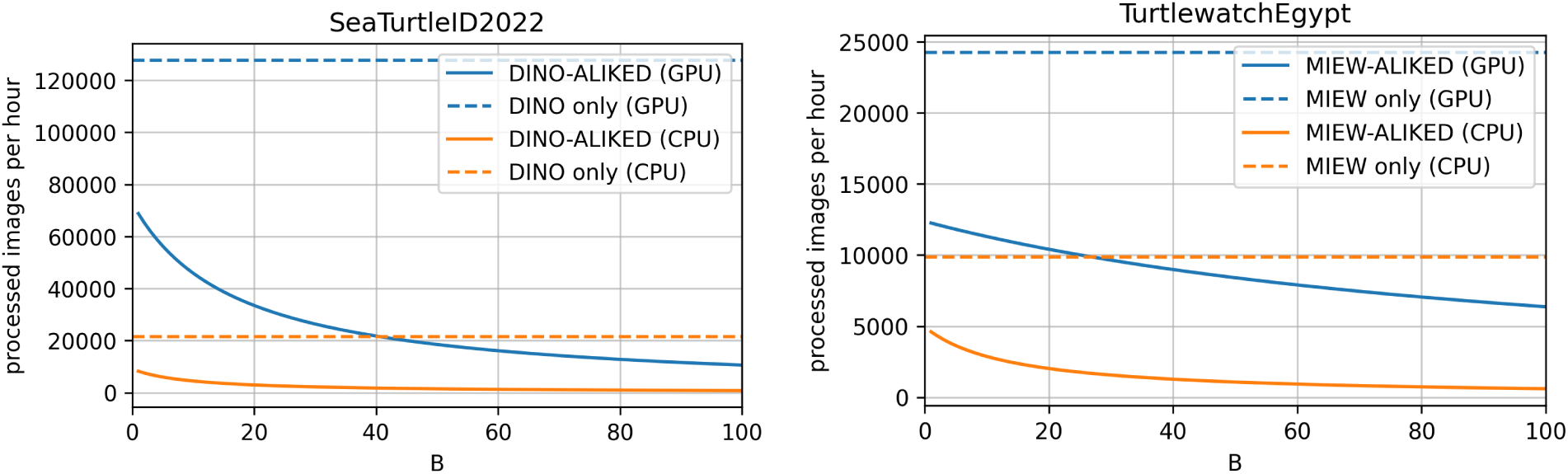
Number of images which can be processed per hour for the pipeline.

## 4. Results

### 4.1. Body parts segmentation results

Figure 6 shows the distribution of the number of body-part segmentations extracted per encounter for the two datasets. Specifically, we report the percentage of encounters from which 0, 1, 2, 3, 4, or at least 5 body-part segmentations were obtained. The distributions are shown separately for each body part (head, front flippers, hind flippers, and full bodies) and for all body parts combined. For the SeaTurtleID2022 dataset (left plot), these distributions simply reflect the availability of the annotated ground-truth segmentations. In contrast, for the TurtlewatchEgypt dataset (right plot), they show the TurtleDetector detections alongside the proposed confidence-based body-part selection protocol.

**Figure 6.**
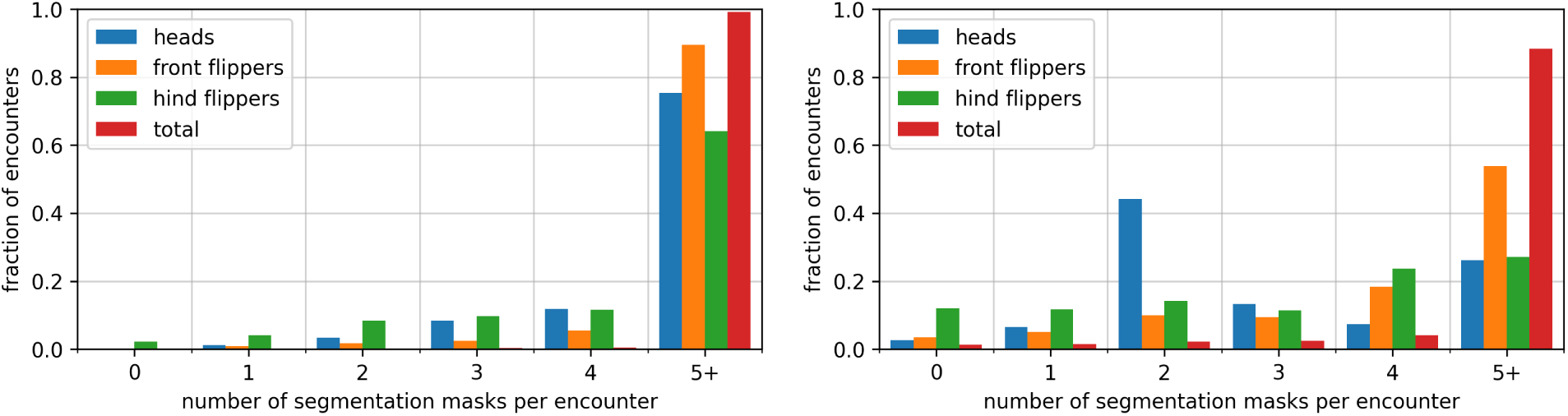
Percentage of encounters from which 0, 1, 2, 3, 4, or at least 5 body-part segmentations were successfully extracted, shown separately for each body part and for all body parts combined. The left plot shows the distribution of ground-truth segmentations for the SeaTurtleID2022 dataset, while the right plot shows the distribution for the Turtlewatch-Egypt dataset obtained by TurtleDetector.

Overall, TurtleDetector performed well on the TurtlewatchEgypt dataset, extracting at least five body-part segmentations in 88.4% of encounters, while no body part was detected in only 1.4% of encounters. Among the individual body parts, hind flippers were extracted least frequently, with no hind-flipper segmentation detected in 12.0% of encounters. We attribute this primarily to the image acquisition setting, as hind flippers are often not visible, for example, when turtles are resting on the seabed, or because of the focus on capturing the heads.

### 4.2. Re-ID results

We first briefly present in the next paragraph, all figures and tables that summarise the results, and then describe them in depth in Sections 4.2.1-4.2.3.

Figure 7 summarises the top-1 accuracies of the main re-ID experiments on the two datasets. As described before, each value represents the average performance across the yearly evaluations. Results are reported for both retrieval strategies: the global-feature model alone (MiewID or DI-NOv3) and the proposed hybrid retrieval framework combining global- and local-feature matching (ALIKED). For the latter, we consider different values of the initial candidate set size *B*. Finally, we report accuracies obtained using each individual body region (heads, front flippers, hind flippers, full bodies) separately, as well as those achieved by the proposed merged multi-body-part approach. Table 4 shows selected top-1 accuracies from Figure 7 for *B* = 10 and 100, while Table 5 shows the corresponding top-5 accuracies for the same experiment. In Figures 8 and 9, we present histograms of the similarity scores corresponding to the image pairs that determine the top-1 predictions. For the single-body-part experiments in Figure 8, the histograms are shown separately for TurtlewatchEgypt and SeaTurtleID2022, and are reported for both retrieval strategies. Figure 9 shows the corresponding histograms for the proposed merged multi-body-part framework and the hybrid retrieval approach only. In these figures, each column represents the body part that ultimately determined the predicted identity by producing the highest similarity score.

**Figure 7.**
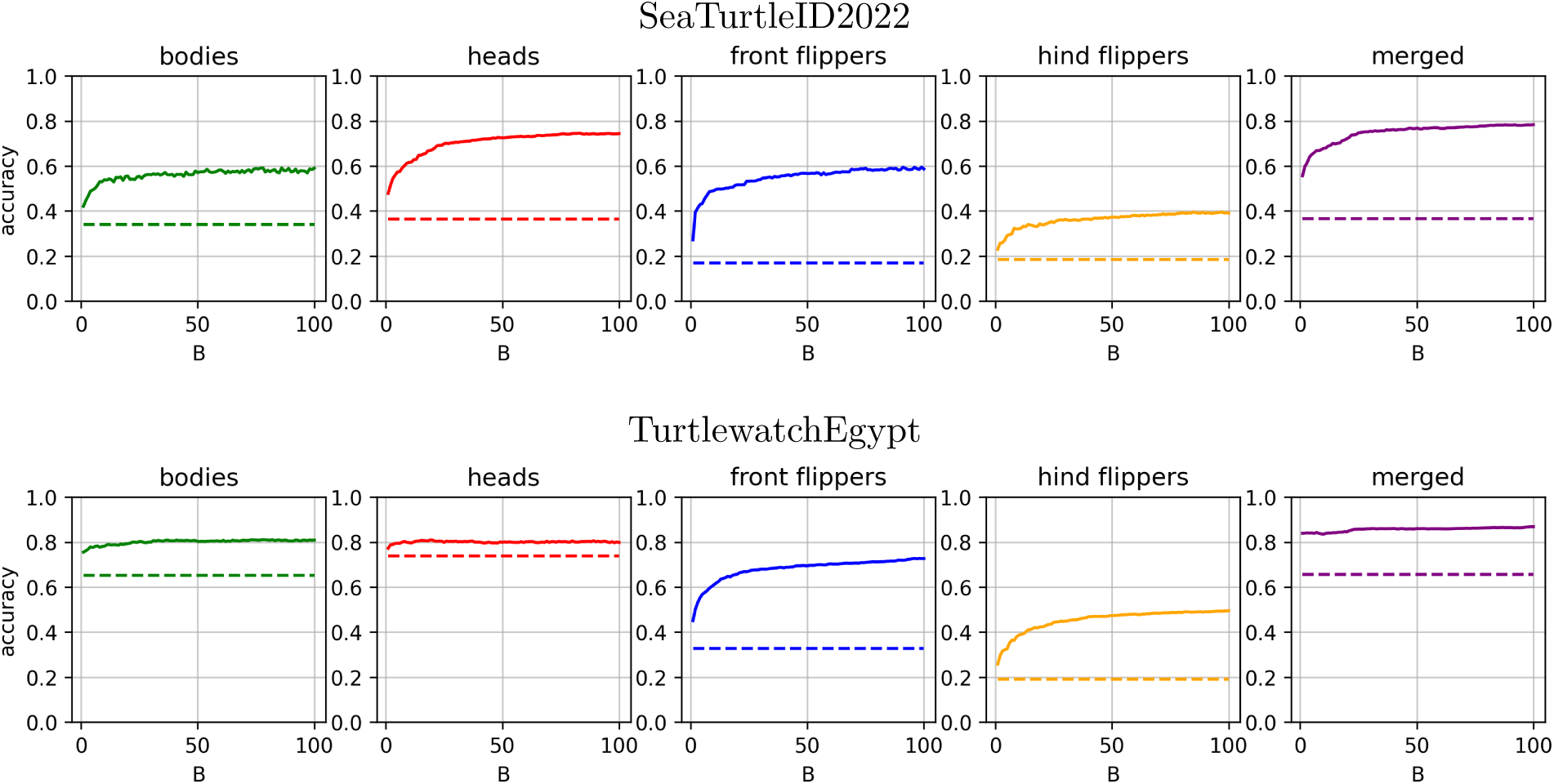
Top-1 accuracies (averaged across years) of the main re-ID experiments for the SeaTurtleID2022 (top) and TurtlewatchEgypt (bottom) datasets. Results are reported for both retrieval strategies (global-feature method only, using MiewID or DINOv3), and the combined global- and local-feature approach (ALIKED) for different values of the budget size *B*. Accuracies are shown separately for each individual single body region used for predictions (heads, front flippers, hind flippers, full bodies) and the proposed merged multi-body-part approach.

**Figure 8.**
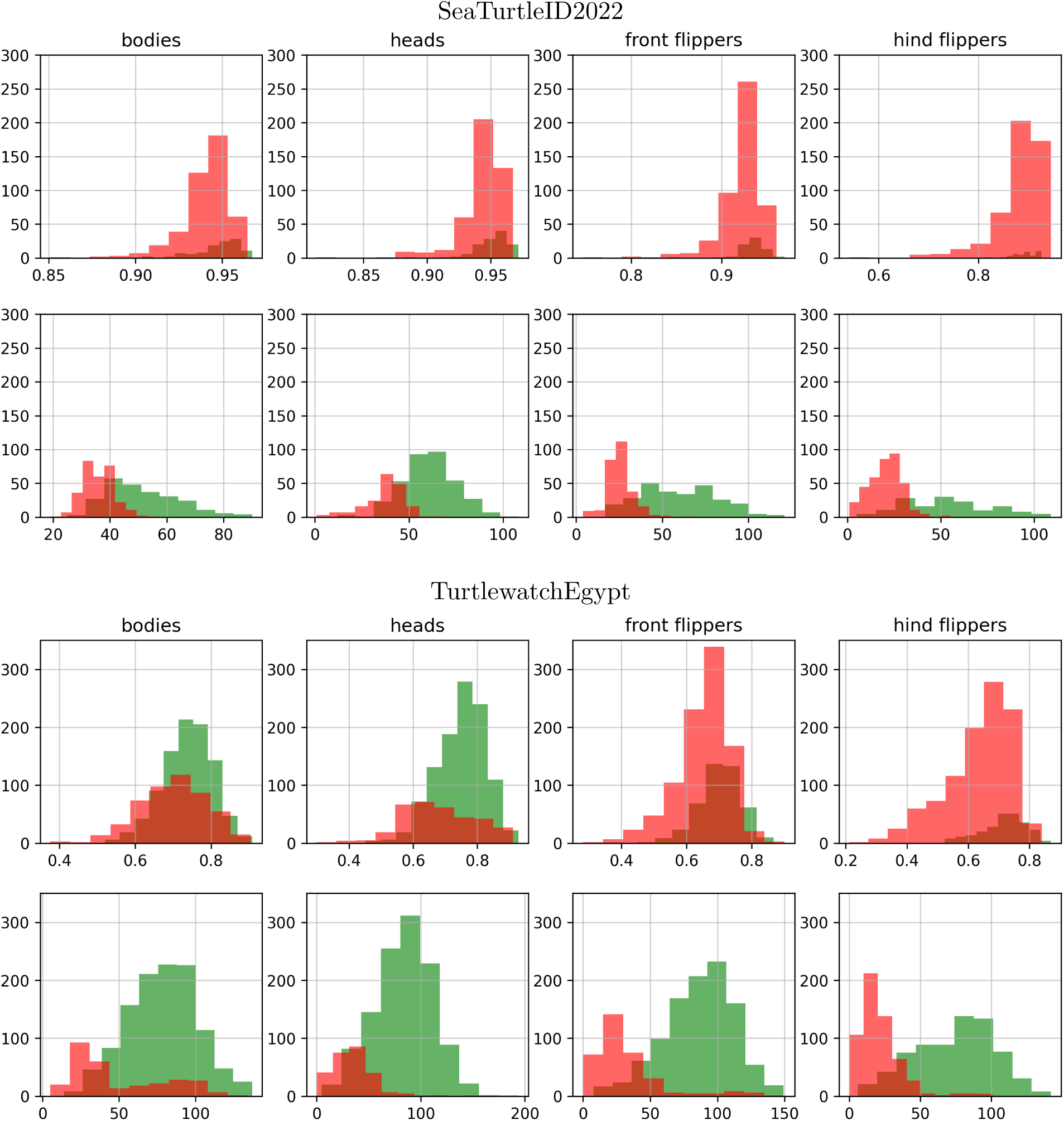
Histograms of the similarity scores corresponding to image pairs that determine the top-1 predictions for the single-body-part experiments for the TurtlewatchEgypt and SeaTurtleID2022 datasets and the two retrieval strategies, MiewID/DINOv3 (top) and MiewID/DINOv3+ALIKED (bottom). Distributions of scores for correct and incorrect matches are shown separately in green and light red, respectively. The overlap between the two distributions is shown in dark red. The area under the red histogram represents the total error.

**Figure 9.**
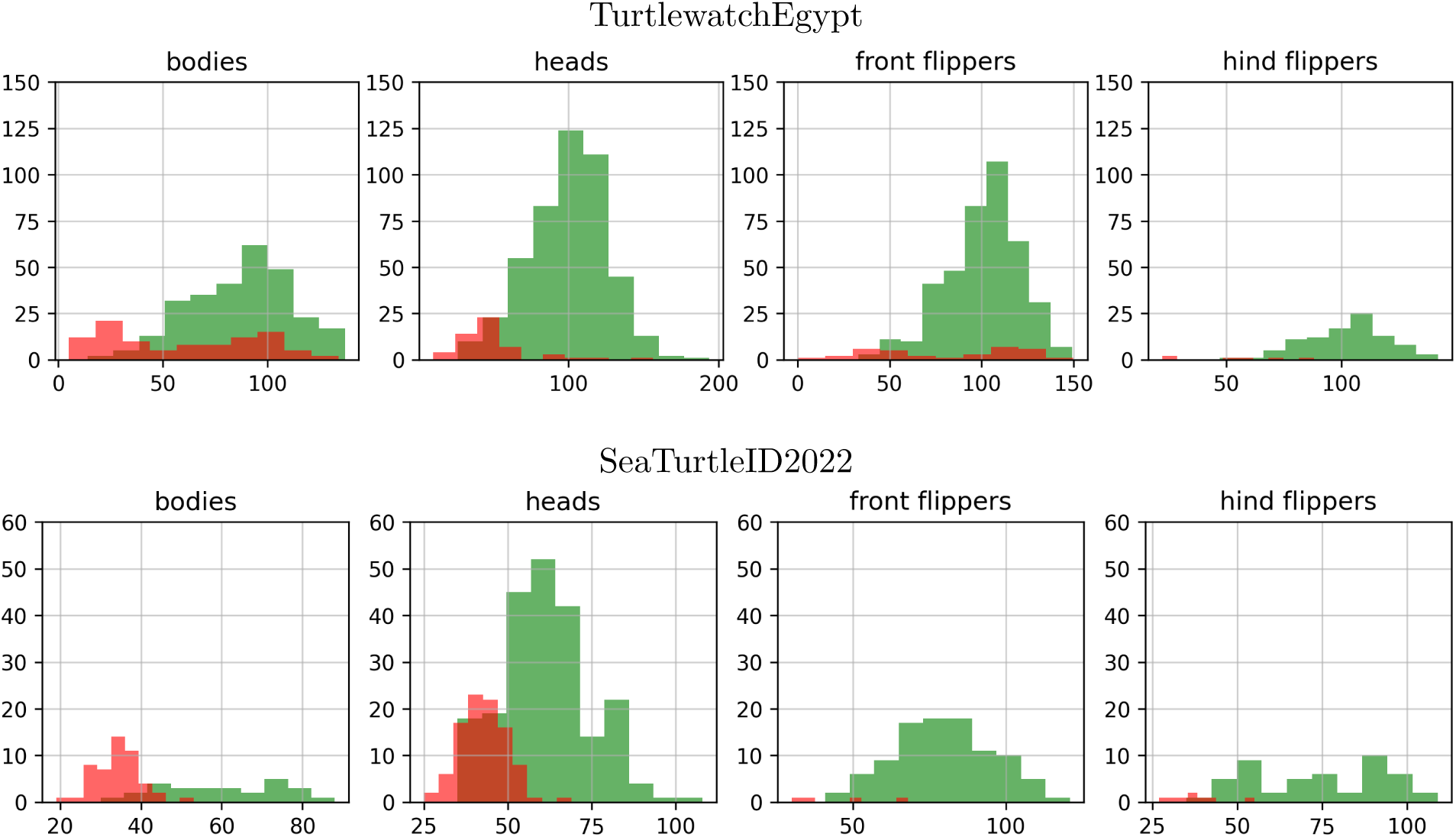
Histograms of the similarity scores corresponding to image pairs that determine the top-1 predictions for the merged-body-part experiments for the TurtlewatchEgypt and SeaTurtleID2022 datasets and the MiewID/DINOv3+ALIKED retrieval strategy. Each column represents the body part that determined the predicted identity by producing the highest similarity score. The colour codes are the same as in Figure 8.

**Table 4.** Top-1 accuracies as values extracted from Figure 7 for *B* ∈ {10, 100}.

| dataset | method | B | bodies | heads | f. flippers | h. flippers | merged |
| --- | --- | --- | --- | --- | --- | --- | --- |
| SeaTurtleID2022 | DINOv3 | - | 34.0 | 36.3 | 16.9 | 18.3 | 36.7 |
|  | DINOv3+ALIKED | 10 | 53.9 | 61.6 | 49.1 | 32.1 | 67.8 |
|  | DINOv3+ALIKED | 100 | 59.1 | 74.4 | 58.7 | 39.1 | 78.4 |
| TurtlewatchEgypt | MiewID | - | 65.3 | 73.9 | 32.8 | 19.2 | 65.7 |
|  | MiewID+ALIKED | 10 | 78.1 | 79.7 | 61.0 | 38.6 | 83.5 |
|  | MiewID+ALIKED | 100 | 80.9 | 79.8 | 72.7 | 49.5 | 86.8 |

**Table 5.** Top-5 accuracies for the same experiment as in Table 4.

| dataset | method | B | bodies | heads | f. flippers | h. flippers | merged |
| --- | --- | --- | --- | --- | --- | --- | --- |
| SeaTurtleID2022 | DINOv3 | - | 54.4 | 55.3 | 40.5 | 34.0 | 55.5 |
|  | DINOv3+ALIKED | 10 | 61.0 | 68.1 | 54.5 | 40.8 | 76.2 |
|  | DINOv3+ALIKED | 100 | 70.3 | 78.7 | 63.6 | 43.7 | 82.3 |
| TurtlewatchEgypt | MiewID | - | 80.2 | 83.3 | 54.8 | 34.3 | 79.1 |
|  | MiewID+ALIKED | 10 | 86.5 | 85.3 | 65.9 | 44.7 | 90.9 |
|  | MiewID+ALIKED | 100 | 88.5 | 84.8 | 76.4 | 53.9 | 92.4 |

#### 4.2.1. Global- versus global- plus local-feature retrieval strategy

From Tables 4 and 5, and Figure 7, we observe that across all experimental settings, incorporating ALIKED (full lines in Figure 7) as the second-stage matcher substantially improves the performance of the global-feature model (dashed lines in Figure 7). For the TurtlewatchEgypt dataset, the improvement is particularly pronounced when identification is based solely on flippers, reflecting MiewID’s relatively poor performance on these body regions. Similarly, DINOv3 achieves comparatively low accuracy on the SeaTurtleID2022 dataset, making the subsequent local-feature refinement by ALIKED especially beneficial. Increasing the budget *B* improves results significantly but saturates relatively quickly.

#### 4.2.2. Single-body-part versus merged multi-body-part framework

Tables 4 and 5, and Figure 7 show that the proposed merged multi-body-part framework consistently outperformed the single-body-part approaches when it was combined with the local matcher ALIKED. Relatively to identification on heads only, the merged approach increased the top-1 accuracy from 74.4% to 78.4% for SeaTurtleID2022 and from 79.8% to 86.8% for TurtlewatchEgypt. In contrast, incorporating multiple body parts was not beneficial when retrieval relied solely on the global-feature model. For the TurtlewatchEgypt dataset, it even significantly reduced performance from 73.9% to 65.7%. We attribute this contrasting behaviour to the separation quality of MiewID and ALIKED shown in Figure 8. While MiewID poorly separates correct and incorrect matches (top row), ALIKED shows a much clearer separation (bottom row). When using MiewID only, high-quality correct matches based on heads, may get overwritten by incorrect matches with high similarity based on flippers. In other words, for MiewID, poor performance on flippers drags down performance in the merged approach, whereas there is a beneficial synergistic effect across multiple body parts for MiewID combined with ALIKED.

#### 4.2.3. Comparison among single-body parts approaches

Considering the single-body-part approaches under the hybrid retrieval strategy, the performance order of body regions was consistent across both datasets. The head achieved most of the times, the highest top-1 accuracy, followed by the full bodies, front flippers, and hind flippers. Although hind flippers were the worst-performing individual body region, they still achieved top-1 accuracies approaching 50% for larger values of *B*, demonstrating that this previously overlooked body region is nevertheless a viable source of identifying information for sea turtle re-ID.

Interestingly, full-body images performed comparably to, and in some cases even better than, head images on the TurtleWatchEgypt dataset, whereas a substantially larger performance gap was observed for the SeaTurtleID2022 dataset. Figure 10 shows representative ALIKED keypoint correspondences for successful top-1 matches across the two datasets. For the TurtlewatchEgypt dataset, which consists predominantly of green turtles, the majority of matched keypoints are located on the carapace, where visible pigmentation patterns are present. In contrast, loggerhead turtles in the SeaTurtleID2022 dataset exhibit few, if any, permanent identifying characteristics on the carapace. Consequently, successful full-body matching relies primarily on the head and flippers. Nevertheless, the accuracy with full bodies is lower than that for these individual body parts. One likely explanation is that these body regions occupy only a fraction of the full-body image. Thus, because global-feature models operate on images of fixed resolution (e.g. 512×512 pixels for MiewID), and we adopt the same fixed resolution for ALIKED, they are represented at a substantially lower resolution than in the corresponding cropped body-part images, reducing the effectiveness of both feature extraction and matching.

**Figure 10.**
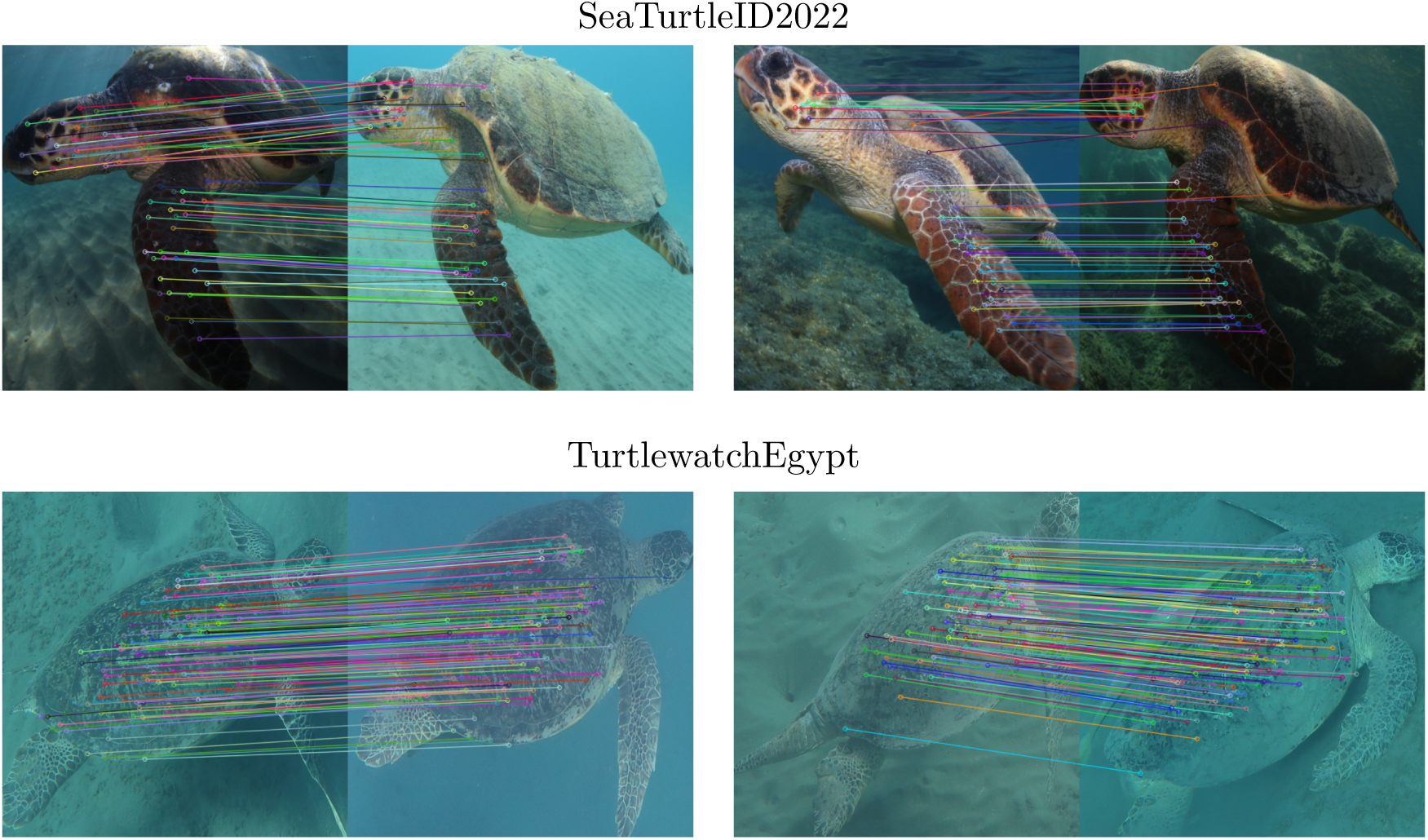
Examples of matching keypoints of ALIKED for successful top-1 predictions for the SeaTurtleID2022 (top) and TurtlewatchEgypt (bottom) datasets when the full bodies are used. Most matches in the latter dataset are based on carapace pigmentation, whereas in the former one, they are predominantly based on heads and flippers. Note the absence of visible carapace pigmentation in the loggerheads from the SeaTurtleID2022 dataset, largely due to algae coverage.

#### 4.2.4. Accuracy gain when incorporating orientation awareness for flippers

In Section 3.5, we introduced orientation-aware flipper matching as one of the strategies in our pipeline, whereby only flippers from the same side are compared. To assess the benefit of this choice, Figure 11 shows the resulting gain in identification accuracy relative to orientation-agnostic matching. We see that indeed there is a benefit from this approach, albeit small; around 1-2% for small values of the budget *B*, but diminishing as the budget grows.

**Figure 11.**
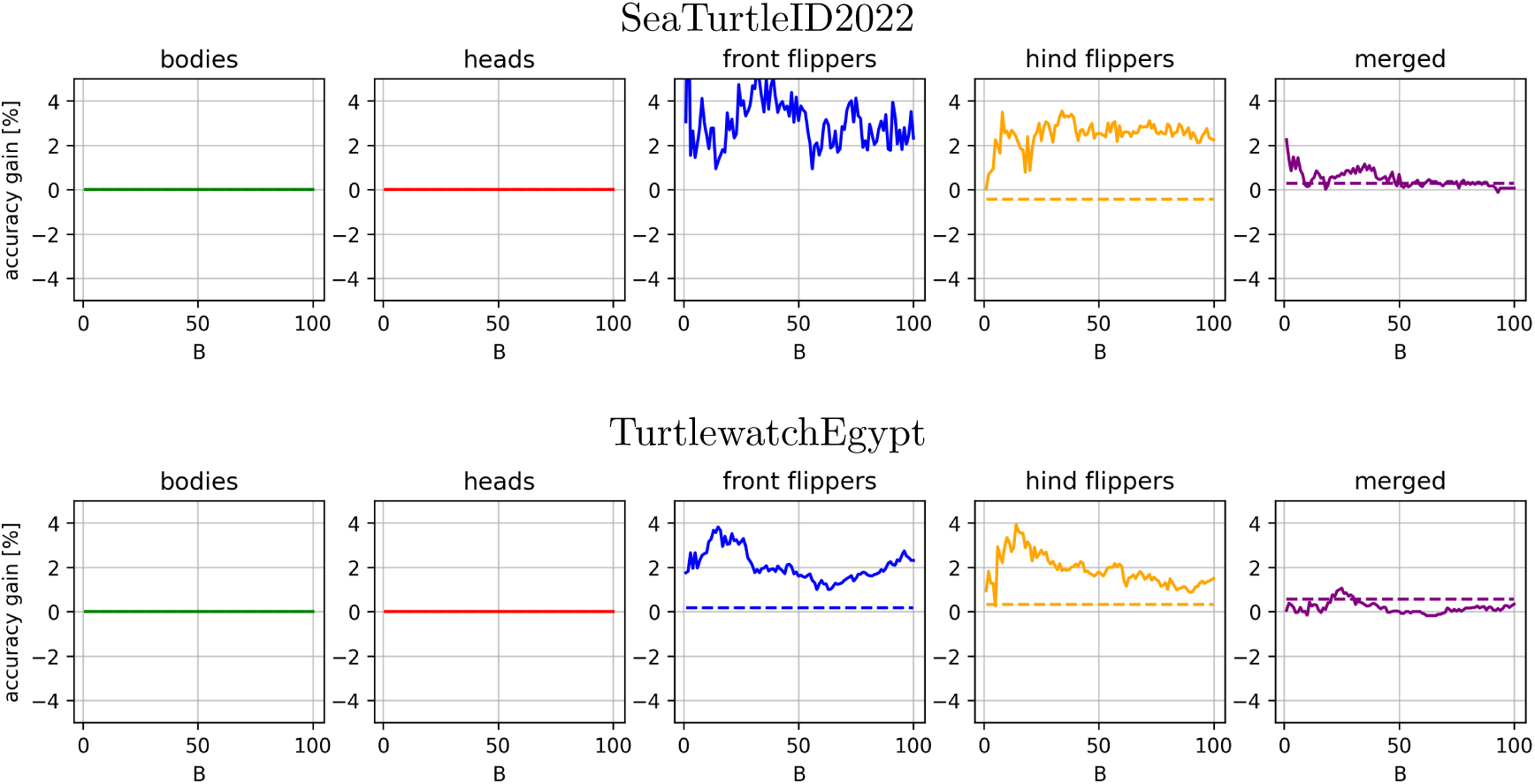
Accuracy gain for the re-ID experiments for SeaTurtleID2022 (top) and Turtle-watchEgypt (bottom) datasets when incorporating orientation awareness for flippers.

## 5. Discussion

In many animal species, multiple body regions exhibit individual-specific patterns that can be exploited for re-ID. However, these complementary identifying patterns are rarely captured in a single image, and the visibility and quality of different body regions often vary across photographs taken during the same encounter. In this study, we introduced an encounter-based, multi-body-part framework that systematically exploits all available identifying information, using sea turtles as a model species. Our experiments demonstrate that integrating information from the head, front flippers, hind flippers and full bodies consistently improves identification performance, yielding a 4–6% increase in top-1 accuracy compared to the best single-body-part approach. These findings suggest that field photo-acquisition protocols could benefit from capturing multiple informative body regions during each encounter, rather than focusing exclusively on a single body part whenever this is practically feasible.

To our knowledge, this is the first study to introduce an encounter-based wildlife re-ID framework that explicitly combines identifying information from multiple body regions. Previous works have largely focused on identifying individuals from a single body region, (Vidal *et al*., 2021; Dunbar *et al*., 2021), on aggregating several images and features of the same body region into a single prediction (Nepovinnykh *et al*., 2024; Nepovinnykh *et al*., 2025) or on fusing complementary similarity measures computed from the same image, such as the combination of global and local features (Cermak *et al*., 2024). In contrast, our approach exploits the complementary identifying information available across multiple photographs and body parts within an encounter, thereby more closely reflecting how ecologists perform manual re-ID in practice.

Among the single-body-part approaches, the highest re-ID accuracy was achieved for heads, followed by full bodies and front flippers, whereas hind flippers yielded the lowest accuracy. We attribute this primarily to the training of current global-feature models, such as MiewID, which have been optimised using head images or full body images rather than flipper segmentations and are therefore better adapted to extracting discriminative features from facial scale patterns (Otarashvili *et al*., 2024; Čermák *et al*., 2024). Consistent with this hypothesis, the performance gap between the head and flippers became substantially smaller when ALIKED was used as the second-stage matcher. As a local-feature method, ALIKED is better suited to exploiting the geometric structure of scale patterns and is therefore less affected by the body region under focus. Note that the strong performance of MiewID on full-body images from the TurtleWatchEgypt dataset, particularly in comparison with the SeaTurtleID2022 dataset, can likely be additionally attributed to the fact that the carapace pigmentation patterns of green turtles resemble those of other species included in MiewID’s training data. A second contributing factor is the image acquisition protocol. Both datasets were originally collected for head-based re-ID, with field efforts focused on obtaining clear images of the facial profiles rather than the flippers or carapaces. Consequently, flippers – particularly the hind flippers – are often only partially visible or captured under less favourable viewpoints and image quality. Despite these limitations, our results demonstrate, for the first time, that reliable sea turtle re-ID based on hind flippers or carapaces (in the absence of algal growth on the carapace) is feasible under unconstrained field conditions. To date, apart from heads, which have been the standard body region for sea turtle re-ID, the literature has explored almost exclusively front-flipper (Pursley, 2020; Mills *et al*., 2023) and, more rarely, carapace identification (Tabuki *et al*., 2021), largely using images acquired under more controlled conditions. Our findings imply that hind flippers and, where informative, the carapace represent valuable, previously underutilised sources of identifying information. Although neither body region alone matches the practicalities and, in some cases, the identification performance of the heads or front flippers, their complementary information can be effectively integrated into the proposed merged multi-body-part re-identification framework, resulting in a consistent net gain in retrieval accuracy.

Although re-ID based solely on other body parts was less accurate than head-based identification, incorporating multiple body-part information into the proposed framework consistently improved overall performance. This demonstrates that the additional identifying information contained in the flippers and carapace complements, rather than merely duplicates, that contained in the head, and highlights the effectiveness of the proposed strategy. Ideally, a multi-body-part identification framework would automatically determine the relative importance of each body part for a given encounter, assigning greater weight to body regions that provide the most reliable identifying information, mimicking the human approach. Despite its simplicity, our strategy of selecting the highest calibrated similarity score across all body parts proved to do exactly that. Nevertheless, more sophisticated approaches could further improve performance. For example, future work could investigate learning body-part calibration weights directly from data or employing fusion strategies that jointly combine evidence from multiple body parts (Cermak *et al*., 2024).

Incorporating multi-body-part information into the final retrieval process was beneficial only when ALIKED was employed as the second-stage matcher. When retrieval relied solely on the global-feature model MiewID, adding this information significantly reduced top-1 accuracy. We attribute this to MiewID’s limited ability to discriminate between flippers of different individuals, often assigning high similarity scores to non-matching flipper pairs as Figure 8 indicates. Consequently, the additional flipper information introduces noise rather than improving retrieval when used within the global-feature model alone, whereas the subsequent use of ALIKED can correct the resulting misrankings. Notably, the combination of global and local feature matching not only exploits the benefits of the merging mechanism but also provides a good compromise between computational efficiency and retrieval accuracy. The number of inexpensive global similarity computations (1) scales linearly with the database size, whereas the number of computationally expensive local comparisons (2) is independent of the database size.

The successful deployment of a multi-body-part re-ID framework relies on the accurate extraction of high-quality body-part detections. We showed that performing re-ID directly on full-body images is generally less effective. From the perspective of local-feature matching, full-body images may contain substantial amounts of irrelevant visual information, increasing the computational cost of descriptor extraction and matching. Conversely, global-feature models operate on images of fixed resolution (e.g. 512×512 pixels for MiewID) and we adopt the same fixed resolution for ALIKED. This means that when the entire animal is resized to the network input resolution, smaller body regions occupy only a small fraction of the image, potentially obscuring fine-scale identifying characteristics. By localising and cropping individual body parts, each region can instead be represented at a much higher effective resolution, preserving the discriminative scale patterns used for identification. These considerations highlight the importance of TurtleDetector, which automatically extracts a small set of high-confidence body-part detections from each image as Figure 6 shows.

TurtleDetector’s ability to predict the orientation of each flipper also proved beneficial. In the flipper-only experiments, orientation-aware matching increased top-1 accuracy by 2–3% by restricting comparisons to the corresponding flipper (e.g. front left flipper to front left flipper), thereby reducing the search space and avoiding unnecessary comparisons between opposite flippers. However, the impact of orientation-aware matching was much smaller in the proposed multi-body-part framework, likely because complementary information from the remaining body parts largely compensated for occasional mismatches. We also note that, in contrast to the flippers, head images were matched in an orientation-independent manner; that is, left and right facial profiles were compared against both left and right profiles in the reference database. Previous work has shown that such an approach is preferable when using global-feature models alone (Adam *et al*., 2025). However, the same study demonstrated that horizontally flipping one of the images before local-feature matching substantially improves the estimation of correspondences between opposite facial profiles. Incorporating this strategy into the proposed hybrid retrieval framework represents another direction for further improving identification performance.

Although our results clearly demonstrate that combining information from multiple body parts improves the robustness of automated re-ID, they do not allow us to fully disentangle potential species-specific differences in the contribution of individual body parts. For example, it has been suggested that flipper-based identification may be particularly advantageous for hawksbill turtles, due to their relatively small number of facial scales and lower inter-individual variability in facial scale patterns compared with other sea turtle species (Papafitsoros *et al*., 2025). However, our experimental design did not permit a rigorous evaluation of this hypothesis. A direct comparison between loggerhead turtles and the other species was not possible because, to avoid data leakage, a different global-feature model was used for the SeaTurtleID2022 dataset than for the TurtlewatchEgypt one. Furthermore, although the TurtlewatchEgypt dataset contains both green and hawksbill turtles, the relatively small number of hawksbill individuals prevented us from treating the two species as separate datasets. Under our evaluation protocol, this would have resulted in an insufficiently large query set for a meaningful comparison. However, the observation that full-body images performed almost as well as head images on the green turtle-dominated TurtleWatchEgypt dataset, in stark contrast to the SeaTurtleID2022 dataset, together with the keypoint correspondences shown in Figure 10, suggests that this difference is largely driven by successful matching of carapace pigmentation. These findings suggest that carapace-based re-ID might be considerably more viable for green turtles than for loggerheads, whose carapaces are frequently obscured by epibionts, reducing the visibility of potentially identifying pigmentation patterns, as was the case in the SeaTurtleID2022 dataset, but not in the TurtlewatchEgypt one. This may partly reflect the greater diversity of epibiont communities reported for loggerheads compared with green turtles (Robinson and Pfaller, 2022). However, future studies using larger, species-specific datasets under diverse acquisition conditions will be needed to determine whether the relative importance of different body parts varies among sea turtle species.

Although demonstrated here using sea turtles, the proposed framework is not species-specific and could be readily extended to other taxa for which multiple body regions provide complementary individual-specific information. For instance, left- and right-side facial or flank asymmetries have long been exploited separately in species such as giraffes, zebras (Parham *et al*., 2017) and lynxes (Picek *et al*., 2026b). Many other species likewise present several body regions suitable for re-ID: Elephants can be identified from ear-notch patterns as well as tusk shapes (Kulits *et al*., 2021); whale sharks and manta rays from flank spot patterns as well as dorsal-fin or ventral markings (Arzoumanian *et al*., 2005; Town *et al*., 2013); and cetaceans from dorsal-fin shape as well as fluke shape and pigmentation (Patton *et al*., 2023), just to name a few examples. In all of these cases, as with sea turtles, field photographs rarely capture all identifying regions in a single frame, and practitioners already manually combine partial evidence when matching individuals. We therefore expect the encounter-based, multi-body-part fusion strategy proposed here to transfer to these settings with comparatively little modification beyond retraining or fine-tuning the detection and matching components for the target species.

## 6. Conclusion

We presented an encounter-based, multi-body-part framework for automated wildlife photo-identification that more closely reflects how ecologists actually perform manual re-ID: By drawing on whichever identifying body regions are visible and well captured across images from an encounter, rather than relying on a single image of a single body part. Applied to sea turtles, the framework consistently outperformed single-body-part approaches, improving top-1 accuracy by 4–6% across three species and two independently curated datasets. Beyond this headline result, our experiments provide the first evidence that hind flippers carry sufficient identifying information for re-ID across loggerhead, green, and hawksbill turtles, and that carapace pigmentation can support reliable re-ID in green turtles specifically – two body regions that had previously received little attention in the sea turtle re-ID literature, largely due to the practical difficulty of manual inspection. More broadly, our results suggest that photo-acquisition protocols in the field will benefit from deliberately capturing multiple body regions per encounter, rather than optimising solely for a single body region, since complementary body parts can compensate when the primary identifying region is occluded, low quality, or simply absent from an encounter. At the same time, although our merging strategy of selecting the single highest calibrated score across body parts, proved effective, its intentional simplicity gives room for further improvement. Thus, future work should explore learned, data-driven body-part weighting, and extensions of the merging mechanism to jointly combine evidence across body parts rather than selecting a single best score.

While we developed and validated the framework on sea turtles, its underlying design, i.e. detecting body parts, followed by body-part-specific matching and merging calibrated scores at the encounter level, is not specific to this taxon. Thus we hope that releasing TurtleDetector and our full pipeline as open source will encourage its adoption and adaptation across other species where identifying patterns are distributed across multiple body regions.

## Footnotes

https://huggingface.co/BVRA/TurtleDetector

